# REV1 Coordinates a Multi-Faceted Tolerance Response to DNA Alkylation Damage and Prevents Chromosome Shattering in *Drosophila melanogaster*

**DOI:** 10.1101/2024.02.13.580051

**Authors:** Varandt Khodaverdian, Tokio Sano, Lara Maggs, Gina Tomarchio, Ana Dias, Connor Clairmont, Mai Tran, Mitch McVey

## Abstract

When replication forks encounter damaged DNA, cells utilize DNA damage tolerance mechanisms to allow replication to proceed. These include translesion synthesis at the fork, postreplication gap filling, and template switching via fork reversal or homologous recombination. The extent to which these different damage tolerance mechanisms are utilized depends on cell, tissue, and developmental context-specific cues, the last two of which are poorly understood. To address this gap, we have investigated damage tolerance responses following alkylation damage in *Drosophila melanogaster*. We report that translesion synthesis, rather than template switching, is the preferred response to alkylation-induced damage in diploid larval tissues. Furthermore, we show that the REV1 protein plays a multi-faceted role in damage tolerance in Drosophila. Drosophila larvae lacking REV1 are hypersensitive to methyl methanesulfonate (MMS) and have highly elevated levels of γ-H2Av foci and chromosome aberrations in MMS-treated tissues. Loss of the REV1 C-terminal domain (CTD), which recruits multiple translesion polymerases to damage sites, sensitizes flies to MMS. In the absence of the REV1 CTD, DNA polymerases eta and zeta become critical for MMS tolerance. In addition, flies lacking REV3, the catalytic subunit of polymerase zeta, require the deoxycytidyl transferase activity of REV1 to tolerate MMS. Together, our results demonstrate that Drosophila prioritize the use of multiple translesion polymerases to tolerate alkylation damage and highlight the critical role of REV1 in the coordination of this response to prevent genome instability.

**Author Summary:** Organisms have evolved several ways to continue copying their DNA when it is damaged, grouped into the categories of translesion synthesis and template switching. These damage tolerance mechanisms prevent replication forks from collapsing when they encounter DNA damage and prevent catastrophic genome instability and cell death. While the proteins and pathways involved in damage tolerance are beginning to be understood at the single cell level, how they are regulated in multicellular organisms is an intriguing question. In this study, we investigated the mechanisms by which Drosophila tolerate alkylation damage during their development. We discovered that tissues containing rapidly dividing diploid cells favor translesion synthesis over template switching, preferentially utilizing different translesion polymerases in a context-dependent manner. Furthermore, we showed that the REV1 protein, best known for its role in recruiting translesion DNA polymerases to damage sites, performs multiple functions during damage tolerance. Together, our results demonstrate that damage tolerance preferences for multicellular organisms may differ from those observed in cultured cells, and establish Drosophila as a useful model system for studying tolerance mechanisms.

## Introduction

Cellular DNA is constantly exposed to both endogenous and exogenous insults, many of which damage the nitrogenous bases. In cells undergoing DNA replication, this base damage can cause replicative polymerases to pause during synthesis, stalling replication forks [1, 2]. Prolonged stalling results in disassembly of the replication machinery and, in severe situations, fork collapse, leading to one-ended DNA double-stranded breaks (DSBs). These breaks are known to promote mutagenesis, chromosome translocations, aberrant recombination, and cell death [3, 4].

To prevent these genome destabilizing events at stalled replication forks, cells have evolved two sets of DNA damage tolerance (DDT) strategies [5]. The first, called template switching, involves the use of error-free homology-directed mechanisms that stabilize stalled forks and prevent their collapse, while allowing for either lesion repair or bypass. Template switching strategies include homologous recombination (HR)-mediated bypass and fork reversal [6-8]. Both DDT mechanisms are stimulated by PCNA K164 polyubiquitylation, which is catalyzed by the Rad5 E3 ubiquitin ligase in budding yeast and its HLTF and SHPRH counterparts in mammals [9-14].

HR-mediated bypass can occur directly at the fork or post-replicatively at single-stranded gaps following repriming [6, 15, 16]. In both cases, the RAD51 protein promotes strand invasion and copying from the recently synthesized nascent lagging strand [17]. Fork reversal, also called fork regression, occurs when DNA translocases and helicases anneal the nascent leading and lagging strands at the fork, forming a 4-way junction often referred to as a “chicken foot” structure [18-20]. Extension of the leading strand using the newly-synthesized lagging strand allows for bypass of the lesion. The regressed fork can then be acted upon by nucleases and helicases to restart replication [21-26]. The regulation of DNA degradation is critical to the success of this mechanism, as uncontrolled nuclease activity at regressed forks has been shown to be detrimental to genome stability [19, 27, 28].

A second type of DDT, called translesion synthesis (TLS), occurs by recruitment of low fidelity TLS polymerases to lesions, enabling damage bypass [29]. TLS can occur ‘on the fly’ at the replication fork, or at single-stranded gaps that result from repriming downstream of the lesion [30]. TLS polymerases include the Y-family polymerases eta (η), iota (ι), kappa (κ), and Rev1, the B-family polymerase zeta (ζ), and the A-family polymerase theta (θ) [31]. Polζ is a multi-subunit enzyme composed of the Rev3 catalytic subunit, two subunits of Rev7, and the Pol31 and Pol32 subunits [32-35]. TLS polymerases have larger active sites that can accommodate damaged bases or mismatches formed between lesions and incoming nucleotides [29]. As a result, TLS polymerases tend to have a lower fidelity than replicative polymerases and are responsible for much of the mutagenesis observed following exposure to DNA damaging agents such as UV and methyl methanesulfonate (MMS) [29, 36].

TLS polymerases are recruited to sites of damage in at least two different ways. In budding yeast, RPA-coated ssDNA accumulates at stalled forks and signals for Rad6 and Rad18 to monoubiquitylate PCNA at lysine 164 [9, 37, 38]. Monoubiquitylated PCNA then recruits TLS polymerases to DNA lesions through interactions with their ubiquitin binding motifs (UBZ in pol η and pol κ, and UBM in pol ι and Rev1) [39]. TLS polymerases can also be recruited to damage sites through interactions with the C-terminal domain (CTD) of Rev1, which uses its BRCT and UBM domains to interact with PCNA at stalled forks and single-stranded gaps [40, 41]. These TLS polymerases can replace the stalled replicative polymerase and insert a nucleotide opposite the damaged base, after which the replicative polymerase resumes synthesis. Depending on the nature of the lesion, TLS polymerases may also act sequentially, with one polymerase responsible for the initial insertion and a second, more processive polymerase extending past the lesion [42-44].

While the involvement of Rev1 in TLS polymerase recruitment is well established, several studies have suggested additional roles for Rev1 in DDT. Unlike other DNA polymerases, Rev1 possesses only deoxycytidyl transferase activity, inserting cytosines opposite DNA damaged guanines and abasic sites [45-47]. Rev1 also functions to promote the bypass of G-quadruplexes and other non-B DNA secondary structures during replication [48, 49]. Furthermore, Rev1 stabilizes Rad51 filaments to prevent degradation of nascent replication tracts in mammalian cells [50], and associates with Rad5 in budding yeast [51, 52].

To date, most studies of DDT have focused on unicellular eukaryotes and immortalized mammalian cell lines. Here, we have investigated DDT in the context of a multicellular organism, *Drosophila melanogaster*, focusing on translesion synthesis and REV1. We find that rapidly dividing diploid tissues in larval Drosophila, but not fly cells growing in culture, rely largely on TLS to tolerate alkylation damage. REV1 plays a multi-faceted role in DDT. While REV1 recruits TLS polymerases via its CTD, in the absence of pol ζ its catalytic activity becomes critically important for DDT. Cells from *rev1* null mutant flies accumulate double strand breaks and experience chromosome shattering when replicating damaged DNA. Interestingly, both Pol η and Pol ζ are used during alkylation damage tolerance, with pol η playing an essential role when TLS is impaired by the deletion of the REV1 CTD. Together, our studies establish Drosophila as a robust genetic system in which to study DNA damage tolerance strategies in a multicellular organism.

## Results

### Drosophila *rev1* mutants are hypersensitive to damaging agents that stall replication forks

We previously showed that *rev1* mutant larvae are sensitive to ionizing radiation (IR) and fail to develop to adulthood post-irradiation [53]. To determine whether this sensitivity is due to a defect in double-strand break repair or an inability to bypass other types of damage created by IR, we tested *rev1* mutant larvae for their ability to survive exposure to other DNA damaging agents. We created a *rev1* null mutant (*rev1Δ*) through imprecise excision of a *P* transposon inserted in the 5’ UTR of the gene. The *rev1Δ* mutants were mildly sensitive to IR, confirming our previous findings (Figure 1A). However, they were not sensitive to topotecan or bleomycin, both of which are known to create DSBs. *rev1Δ* mutants were also sensitive to both nitrogen mustard, which creates intra- and interstrand crosslinks, and hydroxyurea, which depletes dNTP pools. Strikingly, they were hypersensitive to the DNA alkylating agents methyl methanesulfonate (MMS) and ethyl methanesulfonate (EMS), with fewer than 5% of *rev1Δ* homozygotes surviving doses that did not kill heterozygous larvae. Because DNA crosslinks and alkylation damage can lead to stalled replication forks, these results indicate an important role for REV1 during tolerance of fork-blocking lesions.

**Figure 1:**
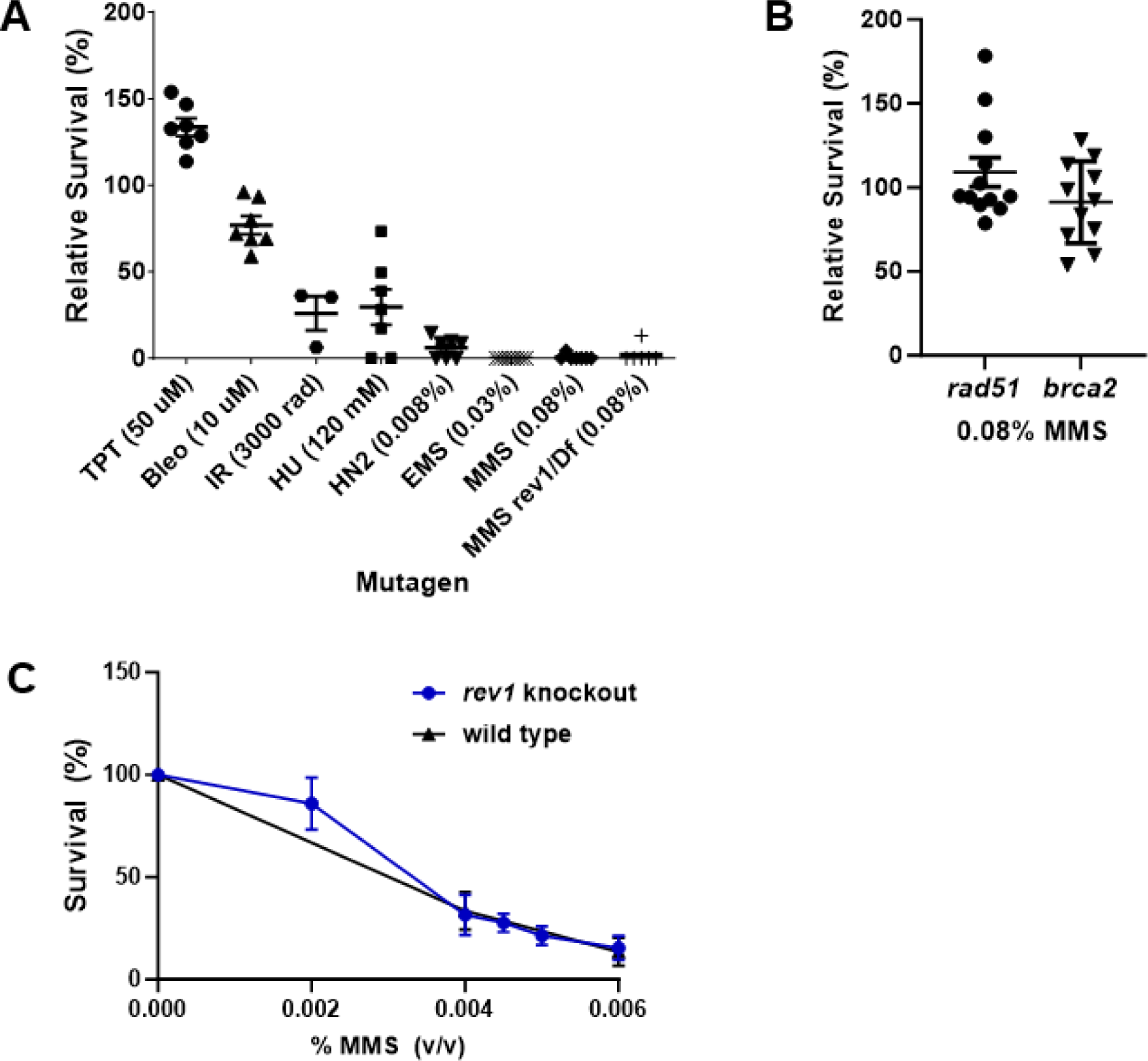
REV1 is vital for tolerance to alkylation damage in *Drosophila melanogaster*, but not S2 cells. (A) Relative survival of homozygous *rev1Δ* mutants to various DNA damaging agents. Heterozygous *rev1Δ* mutants were mated and their larval progeny were treated with indicated concentrations of mutagens or vehicle control in the food. Shown are the percentage of homozygous (or *rev1Δ/ Df(3L)BSC798*) progeny surviving to adulthood, relative to the control. Each point represents one set of control and treated vials, with TTP = topotecan, Bleo = bleomycin, IR = ionizing radiation, HU = hydroxyurea, HN2 = nitrogen mustard, EMS = ethyl methanesulfonate, MMS = methyl methanesulfonate. (B) Relative survival of homozygous *rad51* or *brca2* mutant larvae treated with 0.08% MMS. (C) Survival of wild-type or *rev1* mutant S2 cells treated with increasing concentrations of MMS. Shown are mean and SEM for each genotype.

In *Saccharomyces cerevisiae* and mammalian cells, mutation of genes involved in homologous recombination (HR) repair, such as *RAD51* or *BRCA2*, results in sensitivity to MMS [54-56]. To determine if this is also true in Drosophila, we treated *rad51* and *brca2* null mutants with increasing doses of MMS. These mutants are known to be sensitive to both IR and topotecan [57, 58]. Surprisingly, we observed no sensitivity to a high concentration of MMS in either mutant (Figure 1B). Thus, although HR is critical for repair of double-strand breaks, it is not the primary pathway used to tolerate alkylation damage in Drosophila.

To determine if the *rev1* MMS hypersensitivity is also observed in Drosophila cells grown in culture, we created *rev1* mutant S2 cells via CRISPR-Cas9 genome editing (Supplementary Figure 1). Surprisingly, wild-type and *rev1* mutant S2 cells showed similar sensitivity to increasing concentrations of MMS (Figure 1C), suggesting that unlike flies, immortalized Drosophila cells do not favor TLS for alkylation damage tolerance.

### Loss of REV1 induces double-strand breaks and chromosome aberrations in MMS-treated larval tissues

Larvae treated with lethal doses of DNA damaging agents often survive early development and die prior to pupal eclosion. This is thought to result from massive cell death due to DNA double-strand breaks in rapidly dividing imaginal disc tissues, which are precursors for adult structures including wings, eyes, and other appendages. To test whether this could be responsible for the MMS hypersensitivity observed in *rev1Δ* mutants, we dissected wing imaginal discs from homozygous *rev1Δ* third instar larvae and treated them *ex vivo* with MMS for 5 hours, during which time all cells should replicate their DNA at least once (method per [59]) (Figure 2A). We then quantified the number of γ-H2Av foci, which are indicative of a checkpoint response to double-strand breaks. Strikingly, the number of γ-H2Av foci was 8-fold greater in homozygous *rev1Δ* treated discs compared to heterozygous treated discs (Figure 2B). Together with the survival data, these results indicate that REV1 protects cells in highly proliferative tissues treated with alkylating agents by preventing the formation of double-strand breaks that lead to cell and organismal death.

**Figure 2:**
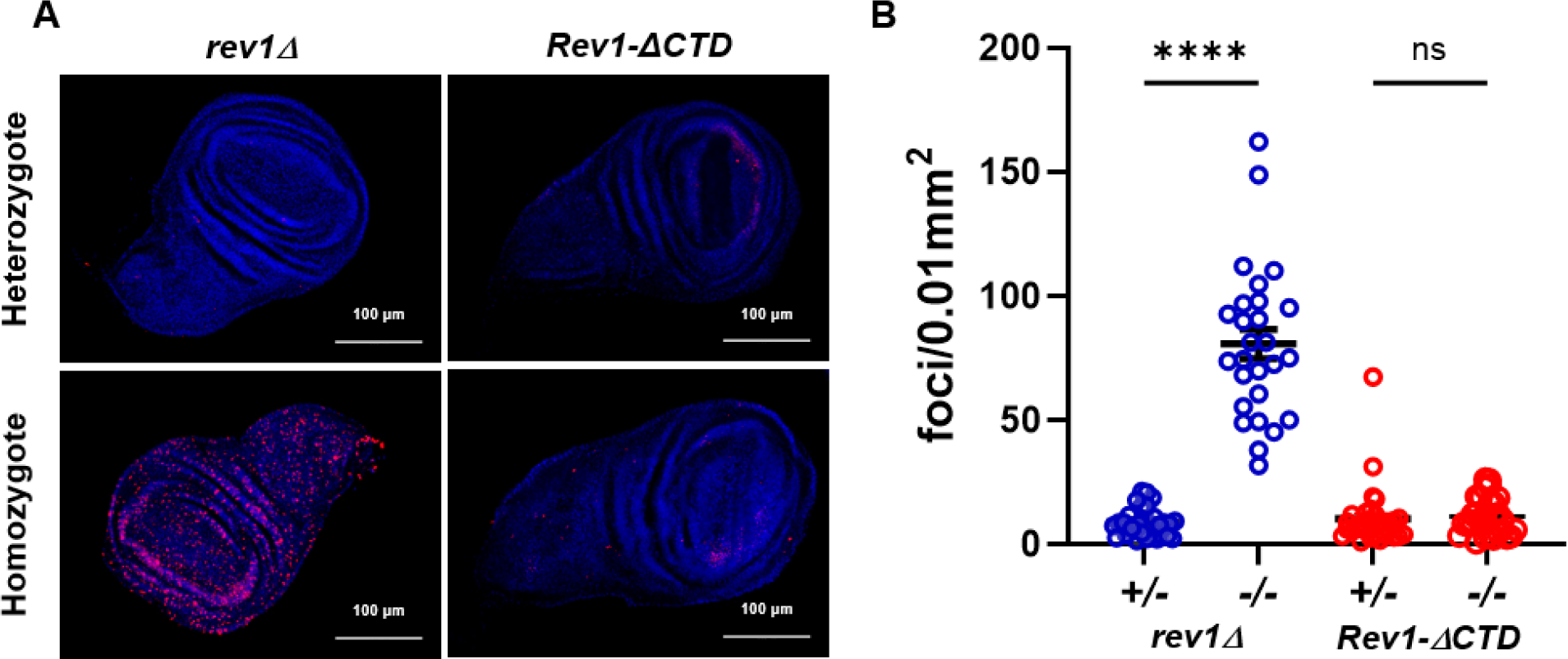
MMS induces DNA double-strand breaks in the absence of REV1. (A) Third instar wing imaginal discs were dissected, treated *ex vivo* for 5 hours with 0.0025% MMS, and stained with DAPI (blue) and an antibody recognizing γ−H2Av (red). Foci were counted and normalized to wing disc size. (B) Quantification of the number of foci in treated discs. Shown are the mean and SEM for each genotype. Statistical comparisons were done using a Kruskal-Wallis one-way ANOVA with Dunn’s multiple comparisons test. n values = 31 (*rev1Δ* +/-), 27 (*rev1Δ* -/-), 31 (*Rev1ΔCTD* +/-), 30 (*Rev1ΔCTD* -/-). **** p<0.0001, ns = not significant.

We wondered whether the increase in double-strand breaks resulting from loss of REV1 might promote chromosome instability. In Drosophila, this instability can be visualized in mitotic spreads from neuroblasts obtained from third instar larval brains. To investigate this question, we dissected brains from wildtype and *rev1Δ* larvae and treated them *ex vivo* for 14 hours with MMS, a period corresponding to approximately two full cell cycles. After obtaining mitotic spreads, we scored them for indicators of chromosome instability, including chromatid breaks and fusions (Figure 3A). While the number of chromatid breaks and chromatid fusions per spread were not significantly different between wild-type and *rev1Δ* flies, we observed a significant increase in a type of catastrophic damage involving chromosome shattering and/or aneuploidy. These events were increased five-fold in *rev1Δ* homozygous neuroblasts treated with MMS, compared to the wild-type control (Figure 3B). These data, combined with the observed increase in γ-H2Av foci in imaginal discs, suggest that upon fork stalling REV1 may prevent the accumulation of double-strand breaks that persist into mitosis and lead to genomic catastrophe.

**Figure 3:**
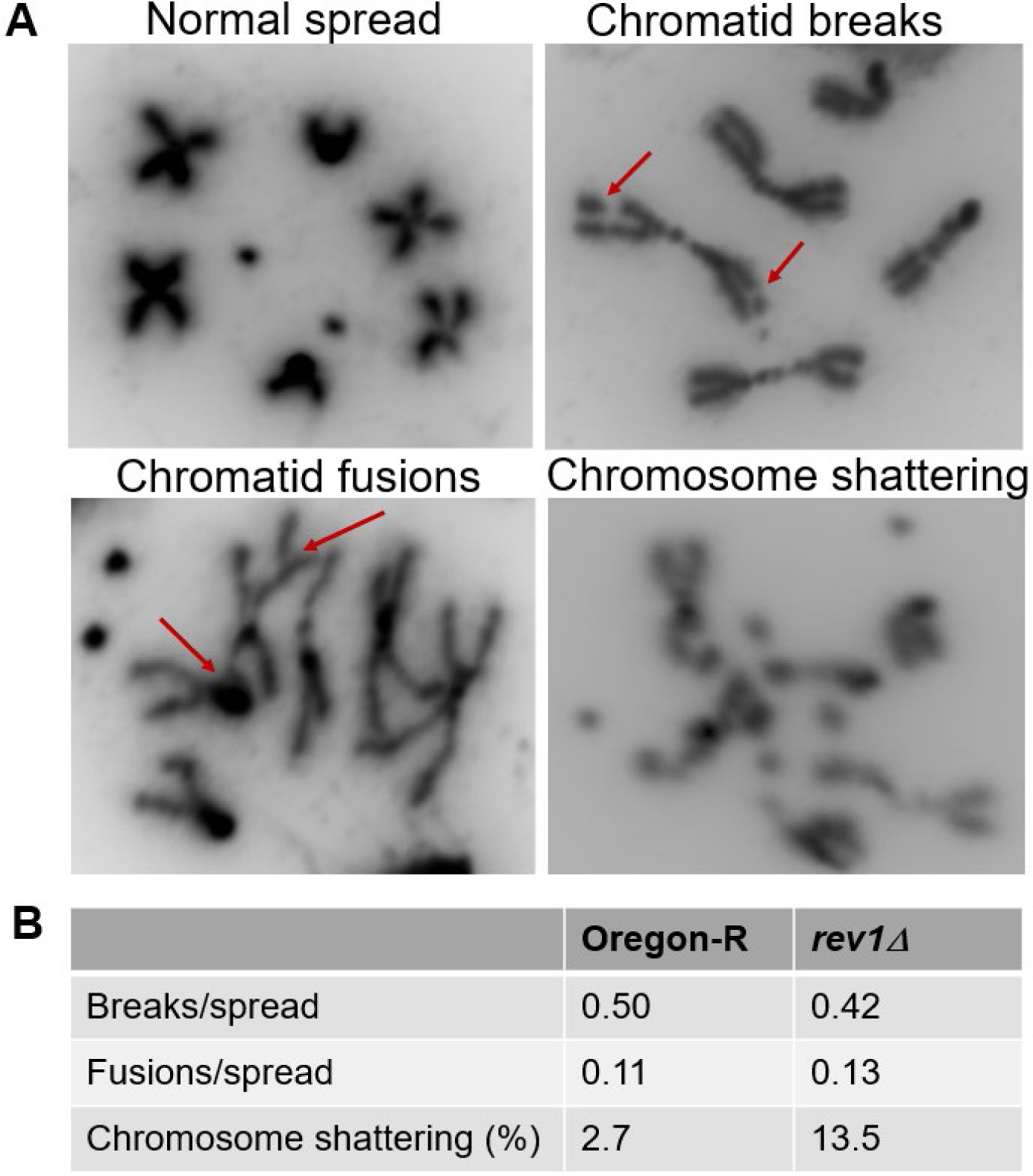
MMS-treated *rev1* neuroblasts have elevated rates of catastrophic chromosome shattering. (A) Representative images of normal and aberrant mitotic spreads. Brains were dissected from wild-type (Oregon-R) and *rev1Δ* third-instar larvae, treated *ex vivo* with 0.0001% (v/v) MMS for 14 hours, incubated with colchicine for 1.5 hours, and squashed. (B) DAPI-stained mitotic spreads were scored for chromatid breaks, chromosome fusions, and catastrophic events (more than three breaks and/or aneuploid spreads). n values = 112 (Oregon-R), 85 (*rev1Δ* -/-).

### Translesion polymerases eta and zeta promote damage tolerance partially independent of the REV1 C-terminal interaction domain

A major role of REV1 in vertebrates is to recruit other translesion polymerases to sites of DNA damage through a physical interaction with its CTD (Figure 4A) [60-64]. This function is conserved in Drosophila, where REV1 interacts with TLS polymerases η, ζ, and ι via its CTD [65]. We previously showed that *rev3Δ* mutants lacking the catalytic subunit of pol ζ are sensitive to alkylating agents [53]. Interestingly, a site-by-side comparison of *rev1Δ* and *rev3Δ* mutants shows that *rev1Δ* mutants are significantly more sensitive to MMS (Figure 4B). This finding differs from observations in *S. cerevisiae*, where *rev1Δ* and *rev3Δ* mutants are equally sensitive to alkylating agents [56].

**Figure 4:**
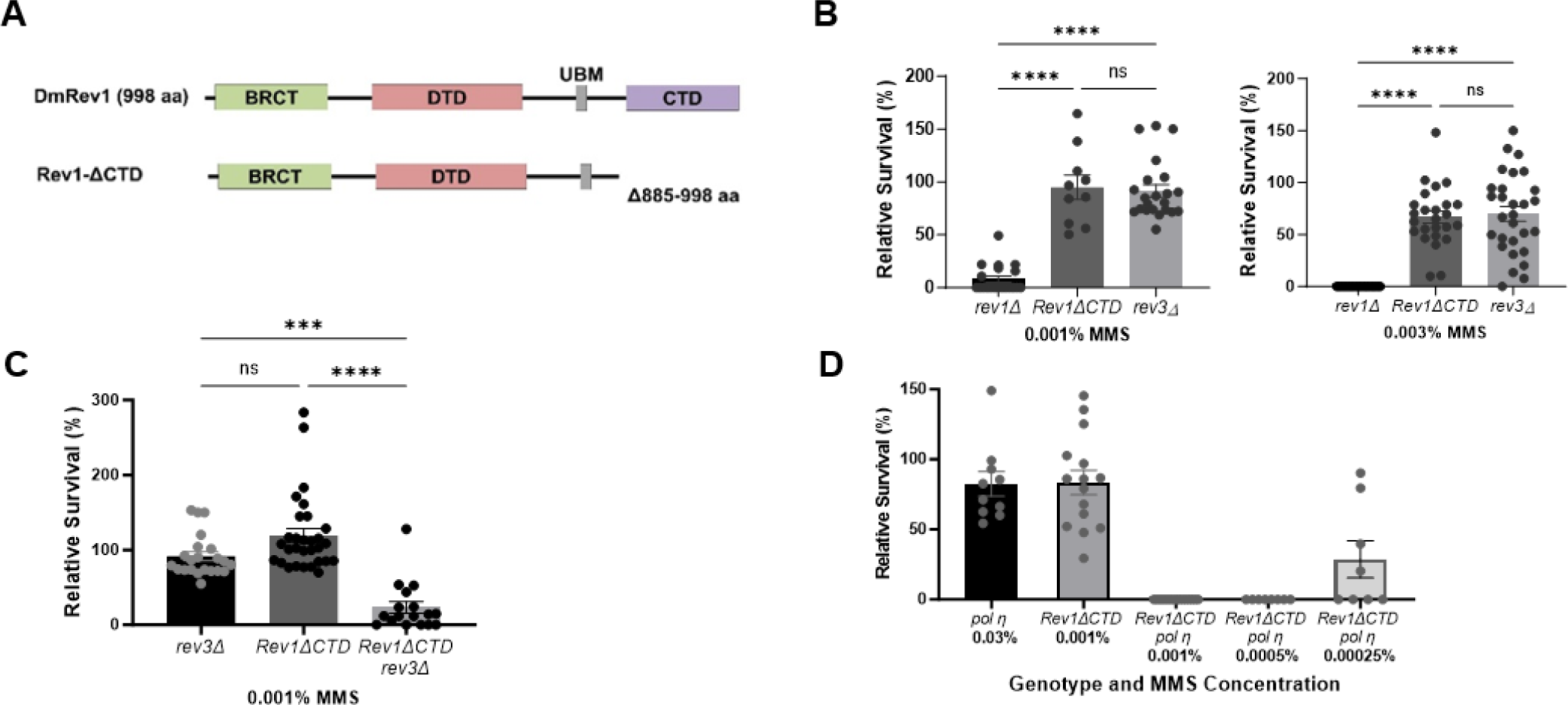
The CTD of REV1 and translesion polymerases η and ζ cooperate to promote MMS-induced damage tolerance. (A) The *Rev1ΔCTD* allele removes the carboxy terminal domain (CTD), which interacts with translesion polymerases η, **ζ**, and ι in Drosophila (Kosarek et al. 2008). (B-D) Relative survival of homozygous DDT mutants to MMS. Heterozygous flies were self-crossed and the resulting larvae were exposed to indicated concentrations of MMS in their food. The percentage of homozygous progeny surviving to adulthood, relative to a vehicle treated control, are indicated. Shown are mean and SEM for each genotype. Statistical comparisons were done using a Kruskal-Wallis one-way ANOVA with Dunn’s multiple comparisons test. * p<0.05, ** p<0.01, *** p<0.001, ****p<0.0001, ns = not significant.

The hypersensitivity of *rev1Δ* mutants could be due to the inability of cells to recruit TLS polymerases to DNA damage sites. Alternatively, REV1 might play another role in damage tolerance. To distinguish between these possibilities, we used site-specific integrase mediated repeated targeting (SIRT) [66-68] to generate an allele of *REV1* lacking the portion of the CTD shown to interact with pol η, pol ζ, and pol ι [65, 69] (Figure 4A). Interestingly, *Rev1ΔCTD* mutants were less sensitive than *rev1Δ* mutants but were equally as sensitive to MMS as *rev3Δ* mutants (Figure 4B). In addition, MMS-treated wing imaginal discs from *Rev1ΔCTD* homozygous mutants did not show increased γ-H2Av foci when compared to heterozygous mutants (Figure 2). Because *rev1Δ* MMS-induced damage and lethality is more severe than that of *Rev1ΔCTD* mutants, we conclude that REV1 plays roles in DDT in addition to TLS polymerase recruitment.

The MMS sensitivity observed in *Rev1ΔCTD* mutants could be due to an inability to recruit one or more translesion polymerases for damage bypass. Y-family polymerases can be recruited to sites of damage through interactions with the Rev1 CTD and through interactions of their UBZ (pol η and pol κ) and UBM (pol ι) domains with monoubiquitylated PCNA [70-72]. To determine if TLS polymerases might also have multiple recruitment mechanisms in Drosophila, we created flies with *Rev1ΔCTD* mutations that were also lacking either REV3 or pol η. Both double mutant stocks were more sensitive to a low concentration of MMS than *Rev1ΔCTD* single mutants (Figure 4C,D). Intriguingly, while loss of pol eta mildly sensitized flies to MMS, the *Rev1ΔCTD pol η* double mutant showed extreme MMS hypersensitivity at doses as low as 0.0025%, suggesting that pol η plays an important role in alkylation damage tolerance when TLS is compromised by loss of the REV1 CTD.

### The deoxycytidyl transferase activity of REV1 becomes important when TLS is compromised

In addition to the CTD, Drosophila REV1 contains a BRCT domain, a deoxycytidyl transferase (DTD) domain, and a single ubiquitin binding motif (UBM) (Figure 5A). In mammals, the BRCT domain interacts with PCNA and with 5’ phosphorylated primer-template junctions [73-75], while the UBM2 domain associates with ubiquitylated PCNA [76, 77]. The DTD catalyzes the insertion of cytosine opposite adducted guanine bases and abasic sites [46, 47]. We used SIRT to create inactivating mutations in each of these domains. The *Rev1ΔBRCT* mutation was created by deleting amino acids 1-121, which corresponds to the entire BRCT domain in mice [78, 79]. The *Rev1-DTD* mutant replaces two amino acids in the catalytic domain with alanines (D421A, G422A), previously shown to abolish deoxycytidyl activity in yeast [80]. Finally, the *Rev1-UBM* mutant changes two conserved residues in the UBM to alanines (L782A, P738A), which impairs the ability of the mouse protein to interact with ubiquitylated PCNA [81]. In all cases, flies with mutations that inactivate each individual domain were not sensitive to MMS (Figure 5B), suggesting that the BRCT, UBM, and DTD domains of REV1 are not required for resistance to MMS-induced damage when TLS is fully functional.

**Figure 5:**
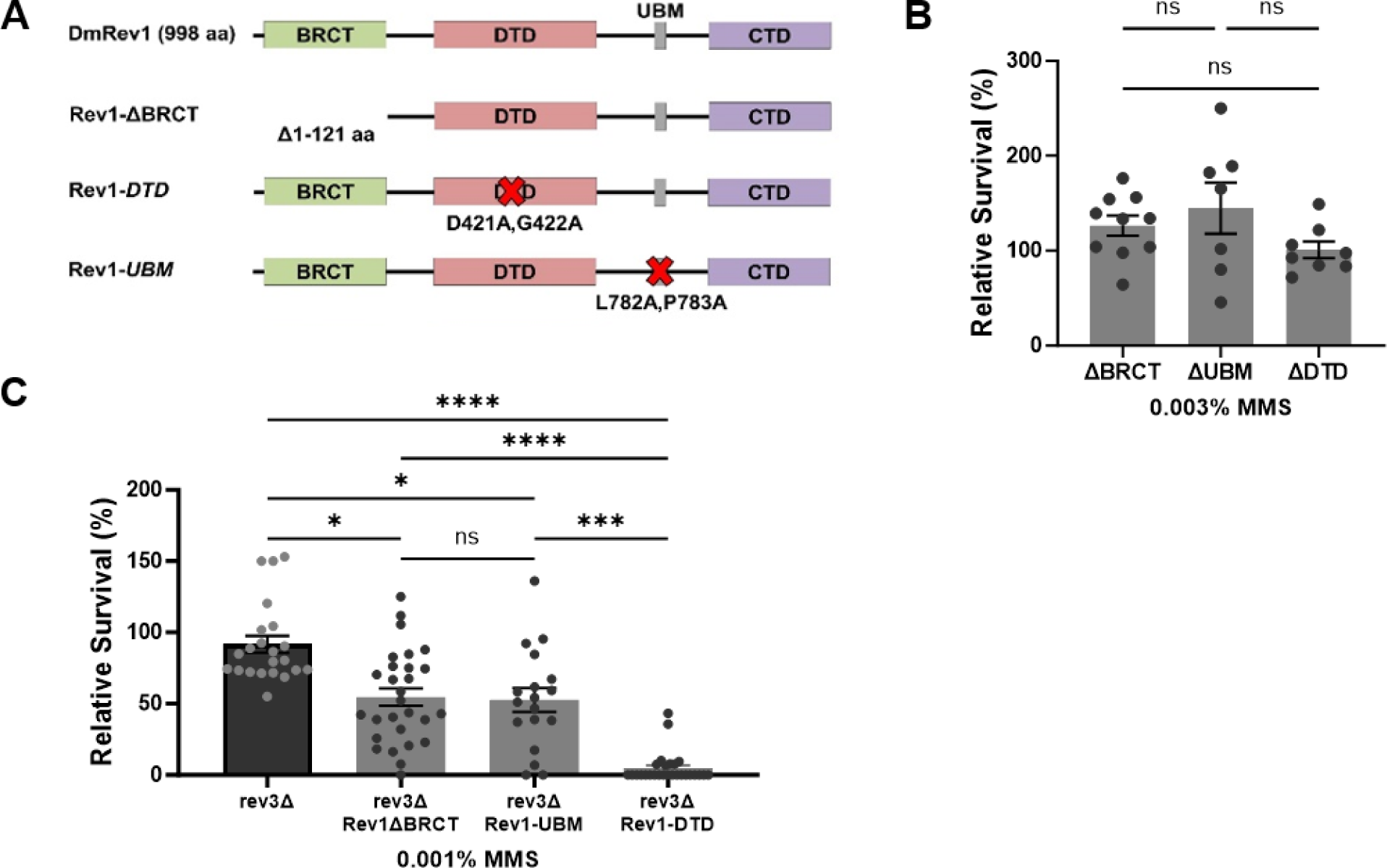
Multiple REV1 domains promote damage tolerance. (A) Domain-specific mutations were created at the endogenous *REV1* locus via SIRT. (B-C) Relative survival of single REV1 domain-specific mutants. Flies heterozygous for the indicated *REV1* domain mutations (B) or homozygous for the *rev3* null mutation and heterozygous for the *REV1* domain-specific mutations (C) were self-crossed and the resulting larvae were exposed to the indicated concentrations of MMS in their food. The percentage of homozygous progeny surviving to adulthood, relative to a vehicle treated control are indicated. Shown are mean and SEM for each genotype. Statistical comparisons were done using a Kruskal-Wallis one-way ANOVA with Dunn’s multiple comparisons test. * p<0.05, *** p<0.001, ****p<0.0001, ns = not significant.

Based on the data shown in Figure 4, the REV3 catalytic subunit of polymerase zeta is important for TLS bypass of MMS-induced damage. To test whether the other domains of REV1 become necessary for MMS resistance when TLS is compromised, we used genetic crosses to place each REV1 domain-specific mutant in a *rev3* background. *Rev1ΔBRCT rev3* and *Rev1-UBM rev3* double mutants were more sensitive to MMS than *rev3* single mutants (Figure 5C). *Rev1-DTD rev3* double mutants showed a greater increase in sensitivity, equivalent to that observed in the *rev1Δ* null mutant (Figure 5C). Based on these data, we speculate that the hypersensitivity of *rev1Δ* mutants to alkylation damage may be largely due to impaired TLS in the absence of the CTD and simultaneous loss of deoxycytidyl transferase activity. In addition, the BRCT and UBM domains may play a minor role in damage tolerance, at least in the absence of Pol ζ.

## DISCUSSION

Here, we have investigated the relative usage of different DNA damage tolerance strategies in Drosophila. Contrary to what has been observed in budding yeast and mammalian cells, homologous-recombination mediated tolerance mechanisms do not appear to be the first line of defense, as *rad51* and *brca2* mutants are not sensitive to high concentrations of MMS. Instead, translesion synthesis, and specifically the REV1 protein, appear to be crucial for damage tolerance. MMS-treated *rev1* mutants accumulate high levels of double-strand breaks in rapidly dividing diploid larval tissues such as imaginal discs. Additionally, chronic exposure to MMS in *rev1* mutants can cause extreme genome instability and chromosome shattering, as we observed in neuroblast mitotic spreads from larval brains. Eventually, these breaks likely lead to extensive cell death, and if compensatory proliferation is unable to restore cell number, cause organismal death prior to adulthood.

Importantly, these studies highlight the importance of studying DNA damage tolerance responses in multicellular organisms, which may show distinct phenotypes compared to cells growing in culture. This is underscored by our observation that *rev1* mutant S2 cells are not hypersensitive to MMS, in contrast to *rev1Δ* mutant flies.

Comparison of the *rev1* MMS sensitivity to that of various domain-specific mutants suggests that REV1 promotes damage tolerance through multiple mechanisms. While *Rev1-ΔCTD* mutants are mildly sensitive to MMS at concentrations of 0.003%, *rev1Δ* mutants cannot survive exposure to 0.001% MMS. Additionally, in contrast to *rev1Δ*, *Rev1ΔCTD* mutant imaginal discs do not accumulate large numbers of double-strand breaks upon MMS exposure. This points to an additional role for Drosophila REV1 in damage tolerance beyond the recruitment of TLS polymerases.

Although the REV1 single domain mutants lacking either BRCT or UBM function are not themselves sensitive to MMS, both showed increased sensitivity to MMS when combined with the loss of the catalytic domain of pol ζ. The BRCT domain has been shown to bind to PCNA for TLS-related functions [73, 74], In addition, in both yeast and mammals the REV1 BRCT domain contains an N-terminal α-helix that can bind to ssDNA, helping recruit it to damage sites [75, 82]. It is possible that the recruitment of REV1 to sites of damage is hindered but not completely abolished without the BRCT domain, due to interactions with ubiquitinated PCNA through the REV1 UBM domain. In humans and yeast, the UBM2 domain of REV1 is responsible for binding with monoubiquitinated PCNA [76, 77]. The similarities in sensitivities between the *Rev1ΔBRCT rev3* and *Rev1-UBM rev3* mutants could correlate with the overlapping functions of these domains in recruiting REV1 to sites of damage. In their absence, Y-family TLS polymerases would be recruited to lesions less effectively, which could result in additive MMS sensitivity when pol ζ is defective.

Interestingly, a greater synergism was seen with the *Rev1-DTD rev3* double mutant. In many contexts, the catalytic activity of REV1 is dispensable. However, the DTD domain can insert cytosine opposite damaged guanines and abasic sites [46, 47, 83]. In Drosophila, without pol ζ, there seems to be a critical role for REV1 deoxycytidyl transferase activity, even when other TLS polymerases are available. It is currently unclear why the DTD domain becomes so important in the absence of pol ζ. Pol ζ is known to be an extender following insertion of a nucleotide opposite a damaged base by Y-family polymerases, but it can also bypass abasic sites on its own [84]. Given the mild sensitivity of pol η mutants, it will be interesting to see if the DTD domain is also critical in the absence of pol η. If so, it may be that the Drosophila REV1 DTD, pol ζ, and pol η have unique but partially overlapping abilities to insert nucleotides opposite different MMS-induced lesions.

One of the interesting results from this study involves the relative MMS resistance of *Rev1ΔCTD* single mutants compared to the hypersensitivity of *Rev1ΔCTD rev3* and *Rev1ΔCTD pol η* double mutants. In chicken DT40 cells, ‘on the fly’ translesion synthesis, which occurs directly at the replication fork, requires the REV1 CTD but not PCNA ubiquitylation [41]. However, post-replicative filling of single-stranded gaps does require PCNA ubiquitylation in DT40 cells. If a similar scenario exists in Drosophila in the *Rev1ΔCTD* mutant, polymerases η and ζ may be compromised in their TLS role at the fork but could still perform their gap-filling functions after fork passage. Loss of either polymerase in a *Rev1ΔCTD* background would compromise TLS bypass both at the fork and during postreplication repair, resulting in enhanced MMS sensitivity. In a REV1 competent background, translesion synthesis bypass at the fork would still be available and could be carried out by polymerases with overlapping abilities, explaining why *pol η* and *pol ζ* mutants are only mildly sensitive to MMS. Validation of this model in Drosophila will require experiments in a genetic background in which gap filling by TLS polymerases is compromised, as might occur in a PCNA K164R mutant that is unable to be ubiquitylated.

We have shown that the BRCT, UBM, DTD, and CTD domains of REV1 all play roles in DNA damage tolerance. Due to the extreme sensitivity of *rev1Δ* mutants, it is possible that other REV1 protein regions are also important. For example, REV1 could be important for stabilizing regressed forks, recruitment of proteins important for fork reversal or template switching, protection of regressed forks from cleavage by structure-specific endonucleases, and/or prevention of hyper-resection. We are currently investigating these possibilities.

Notably, these studies may have relevance to cancer research, as mutagenic TLS is strongly implicated in carcinogenesis, tumor progression, and chemotherapeutic resistance [85-87]. Pertinent to this study, suppression of Rev1 is known to inhibit both cisplatin- and cyclophosphamide-induced mutagenesis, which sensitizes tumors to traditional therapeutics and suppresses the development of tumor chemoresistance [88]. A novel small molecule, JH-RE-06, induces REV1 dimerization and inhibits TLS, making it attractive as a potential therapeutic [89, 90]. As we have shown that TLS is a vital damage tolerance mechanism in Drosophila, we propose this model system may be useful for studying strategies employed by tumor cells exposed to fork-stalling agents and inhibitors of these processes.

## Acknowledgments

We thank Alice Witsell and Ethan Brown for creating the *rev1* mutant S2 cells. This work was supported by NIGMS grant R01GM125827 to M. McVey.

## Materials and Methods

### Drosophila husbandry and stocks

Flies were raised on standard cornmeal agar at 25°C on a 12 hr:12 hr light/dark cycle. The *rev1Δ* null allele was generated by an imprecise excision screen using *P[48]Rev1[G18538]* (Bloomington Stock #28417) with the *P* element inserted 55 bp of the activation start site for *REV1*. The imprecise excision deletes 4531 bp downstream of the *P* element (the entire *REV1* gene), with 951 bp of the *P* element remaining. The *pol η* and *rev3* (encoded by *mus205*) knockout alleles were generated previously in the lab through imprecise *P*-element excision [53]. The *rad51* (encoded by *spn-A*) mutants were compound heterozygotes of *spn-A^093A^* and *spn-A^057^* [91]. The *brca2^KO^*null allele replaces the entire coding sequence of the *BRCA2* gene with the *mini-white* gene [92].

### Endogenous *Rev1* mutant generation

Endogenous *REV1* domain mutants were generated through site-specific integrase mediated repeated targeting (SIRT) [66, 67]. The *Rev1-ΔBRCT* allele deletes the first 121 amino acids of REV1. *Rev1-DTD* is a double D421A, G422A mutation within the catalytic domain of REV1. *Rev1-UBM* is a double L782A, P738A mutation within the conserved region of the UBM domain. The *Rev1ΔCTD* allele deletes the last 113 amino acids (885-998) of REV1.

### Mutagen sensitivity assays

Heterozygous mutant males and females were mated by placing them in a vial for three days, then placed into another set of vials for three more days before being removed. The first set of vials were treated with 250 μL of mutagen, diluted in ddH_2_O, the day after the parental flies were removed (treatment vials). The second set of vials were treated with 250 μL of ddH_2_O the day after the parental flies were removed (vehicle control vials). The number of homozygous and heterozygous eclosed flies were counted in control and treated vials. The relative survival for each vial was calculated as the percent of homozygotes in the treated vials divided by the percent of homozygotes in the control vials.

### Imaginal disc culture and immunofluorescence

Third instar wing imaginal discs were dissected and cultured for 5 hours at 25°C in 20% fetal bovine serum (FBS), 0.7% sodium chloride, 0.1% dimethyl sulfoxide (DMSO) and 0.0025% MMS. Following 5 hours of culture, 90% of wing imaginal disc cells have entered S-phase [93]. Discs were washed twice with cold phosphate buffered saline with 0.1% Tween 20 (1xPBST), fixed with formaldehyde, and incubated overnight at 4°C in 1:500 anti-γH2Av antibody (Rockland Inc.) in 5% bovine serum albumin (BSA) in 1xPBS containing 0.3% Triton X-100. Discs were washed 4X for 5 minutes with PBST and incubated for 2 hours at room temperature in 1:1000 goat anti-Rabbit IgG Rhodamine Red conjugated antibody and 500 μg/mL DAPI in 1xPBS + 5% BSA. Discs were washed and mounted in VECTASHEILD on microscope slides [59]. γ-H2Av foci were imaged at 10-20x magnification using a Zeiss Z-stacking microscope and with filter sets compatible with DAPI and Rhodamine. Discs were imaged multiple times along the Z-axis, processed by deconvolution, and compressed into one image by extended depth of field algorithms. The area of the disc and number of foci per disc were calculated using ImageJ.

### Mitotic chromatid spreads

Incubation of third instar larval brains were modified from Gatti and colleagues [94]. Third instar larval brains were dissected and cultured in 20% FBS, 0.7% sodium chloride, 0.1% DMSO and 0.001% MMS for 14 hours at 25 °C. Colchicine was added to a concentration of 50 μM and the discs were cultured for an additional 1.5 hours. Larval brains were swollen by incubating for 10 minutes in 0.5% sodium citrate, fixed for 20 seconds in Acetic acid, methanol, and pico-pure H2O (5.5:5.5:1), and placed into a drop of 45% Acetic acid on siliconized coverslips. Poly-L-lysine coated slides were placed onto the coverslip and pressure was gently applied for 10 seconds. Complete spreading of mitotic chromatids was achieved by squishing the coverslip and slide using a clamp. Slides and coverslips were then frozen for 15 minutes at -80°C. Coverslips were removed and slides placed into -20°C ethanol for 20 minutes. Slides were removed from the ethanol and dried vertically at room temperature or overnight at 4°C. Slides were rehydrated in 2xSCC for 5 minutes at room temperature. Slides were then incubated for 5 minutes in 2xSCC with 200 μg/mL DAPI for 5 minutes. Slides were washed twice for 10 seconds with 2xSCC, dried at room temperature, and then mounted using Vectashield. Mitotic chromatid spreads were imaged at 100x magnification with a DAPI filter set. Scoring of chromosome aberrations was conducted with blinding, with each spread scored by 2 individuals.

## References

1. Eichman BF. Repair and tolerance of DNA damage at the replication fork: A structural perspective. Curr Opin Struct Biol. 2023;81:102618. Epub 20230601. doi: 10.1016/j.sbi.2023.102618. PubMed PMID: 37269798; PubMed Central PMCID: PMCPMC10525001.

2. Tubbs A, Nussenzweig A. Endogenous DNA Damage as a Source of Genomic Instability in Cancer. Cell. 2017;168(4):644–56. doi: 10.1016/j.cell.2017.01.002. PubMed PMID: 28187286; PubMed Central PMCID: PMCPMC6591730.

3. Zeman MK, Cimprich KA. Causes and consequences of replication stress. Nat Cell Biol. 2014;16(1):2–9. doi: 10.1038/ncb2897. PubMed PMID: 24366029; PubMed Central PMCID: PMCPMC4354890.

4. Pilzecker B, Buoninfante OA, Jacobs H. DNA damage tolerance in stem cells, ageing, mutagenesis, disease and cancer therapy. Nucleic Acids Res. 2019;47(14):7163–81. doi: 10.1093/nar/gkz531. PubMed PMID: 31251805; PubMed Central PMCID: PMCPMC6698745.

5. Marians KJ. Lesion Bypass and the Reactivation of Stalled Replication Forks. Annu Rev Biochem. 2018;87:217–38. Epub 20180103. doi: 10.1146/annurev-biochem-062917-011921. PubMed PMID: 29298091; PubMed Central PMCID: PMCPMC6419508.

6. Branzei D, Psakhye I. DNA damage tolerance. Curr Opin Cell Biol. 2016;40:137–44. Epub 20160406. doi: 10.1016/j.ceb.2016.03.015. PubMed PMID: 27060551.

7. Branzei D, Szakal B. DNA damage tolerance by recombination: Molecular pathways and DNA structures. DNA Repair (Amst). 2016;44:68–75. Epub 20160516. doi: 10.1016/j.dnarep.2016.05.008. PubMed PMID: 27236213; PubMed Central PMCID: PMCPMC4962778.

8. Neelsen KJ, Lopes M. Replication fork reversal in eukaryotes: from dead end to dynamic response. Nat Rev Mol Cell Biol. 2015;16(4):207–20. Epub 20150225. doi: 10.1038/nrm3935. PubMed PMID: 25714681.

9. Hoege C, Pfander B, Moldovan GL, Pyrowolakis G, Jentsch S. RAD6-dependent DNA repair is linked to modification of PCNA by ubiquitin and SUMO. Nature. 2002;419(6903):135-41. doi: 10.1038/nature00991. PubMed PMID: 12226657.

10. Motegi A, Sood R, Moinova H, Markowitz SD, Liu PP, Myung K. Human SHPRH suppresses genomic instability through proliferating cell nuclear antigen polyubiquitination. J Cell Biol. 2006;175(5):703–8. Epub 20061127. doi: 10.1083/jcb.200606145. PubMed PMID: 17130289; PubMed Central PMCID: PMCPMC2064669.

11. Motegi A, Liaw HJ, Lee KY, Roest HP, Maas A, Wu X, et al. Polyubiquitination of proliferating cell nuclear antigen by HLTF and SHPRH prevents genomic instability from stalled replication forks. Proc Natl Acad Sci U S A. 2008;105(34):12411–6. Epub 20080821. doi: 10.1073/pnas.0805685105. PubMed PMID: 18719106; PubMed Central PMCID: PMCPMC2518831.

12. Unk I, Hajdu I, Fatyol K, Szakal B, Blastyak A, Bermudez V, et al. Human SHPRH is a ubiquitin ligase for Mms2-Ubc13-dependent polyubiquitylation of proliferating cell nuclear antigen. Proc Natl Acad Sci U S A. 2006;103(48):18107–12. Epub 20061115. doi: 10.1073/pnas.0608595103. PubMed PMID: 17108083; PubMed Central PMCID: PMCPMC1838714.

13. Unk I, Hajdu I, Fatyol K, Hurwitz J, Yoon JH, Prakash L, et al. Human HLTF functions as a ubiquitin ligase for proliferating cell nuclear antigen polyubiquitination. Proc Natl Acad Sci U S A. 2008;105(10):3768–73. Epub 20080303. doi: 10.1073/pnas.0800563105. PubMed PMID: 18316726; PubMed Central PMCID: PMCPMC2268824.

14. Unk I, Hajdu I, Blastyak A, Haracska L. Role of yeast Rad5 and its human orthologs, HLTF and SHPRH in DNA damage tolerance. DNA Repair (Amst). 2010;9(3):257–67. Epub 20100121. doi: 10.1016/j.dnarep.2009.12.013. PubMed PMID: 20096653.

15. Prado F. Homologous Recombination: To Fork and Beyond. Genes (Basel). 2018;9(12). Epub 20181204. doi: 10.3390/genes9120603. PubMed PMID: 30518053; PubMed Central PMCID: PMCPMC6316604.

16. Khatib JB, Nicolae CM, Moldovan GL. Role of Translesion DNA Synthesis in the Metabolism of Replication-associated Nascent Strand Gaps. J Mol Biol. 2024;436(1):168275. Epub 20230913. doi: 10.1016/j.jmb.2023.168275. PubMed PMID: 37714300.

17. Liu W, Saito Y, Jackson J, Bhowmick R, Kanemaki MT, Vindigni A, Cortez D. RAD51 bypasses the CMG helicase to promote replication fork reversal. Science. 2023;380(6643):382-7. Epub 20230427. doi: 10.1126/science.add7328. PubMed PMID: 37104614; PubMed Central PMCID: PMCPMC10302453.

18. Zellweger R, Dalcher D, Mutreja K, Berti M, Schmid JA, Herrador R, et al. Rad51-mediated replication fork reversal is a global response to genotoxic treatments in human cells. J Cell Biol. 2015;208(5):563–79. doi: 10.1083/jcb.201406099. PubMed PMID: 25733714; PubMed Central PMCID: PMCPMC4347635.

19. Qiu S, Jiang G, Cao L, Huang J. Replication Fork Reversal and Protection. Front Cell Dev Biol. 2021;9:670392. Epub 20210510. doi: 10.3389/fcell.2021.670392. PubMed PMID: 34041245; PubMed Central PMCID: PMCPMC8141627.

20. Ito M, Fujita Y, Shinohara A. Positive and negative regulators of RAD51/DMC1 in homologous recombination and DNA replication. DNA Repair (Amst). 2024;134:103613. Epub 20231213. doi: 10.1016/j.dnarep.2023.103613. PubMed PMID: 38142595.

21. Delamarre A, Barthe A, de la Roche Saint-Andre C, Luciano P, Forey R, Padioleau I, et al. MRX Increases Chromatin Accessibility at Stalled Replication Forks to Promote Nascent DNA Resection and Cohesin Loading. Mol Cell. 2020;77(2):395–410 e3. Epub 20191120. doi: 10.1016/j.molcel.2019.10.029. PubMed PMID: 31759824.

22. Berti M, Ray Chaudhuri A, Thangavel S, Gomathinayagam S, Kenig S, Vujanovic M, et al. Human RECQ1 promotes restart of replication forks reversed by DNA topoisomerase I inhibition. Nat Struct Mol Biol. 2013;20(3):347–54. Epub 20130210. doi: 10.1038/nsmb.2501. PubMed PMID: 23396353; PubMed Central PMCID: PMCPMC3897332.

23. Conti BA, Smogorzewska A. Mechanisms of direct replication restart at stressed replisomes. DNA Repair (Amst). 2020;95:102947. Epub 20200816. doi: 10.1016/j.dnarep.2020.102947. PubMed PMID: 32853827; PubMed Central PMCID: PMCPMC7669714.

24. Pasero P, Vindigni A. Nucleases Acting at Stalled Forks: How to Reboot the Replication Program with a Few Shortcuts. Annu Rev Genet. 2017;51:477–99. doi: 10.1146/annurev-genet-120116-024745. PubMed PMID: 29178820.

25. Thangavel S, Berti M, Levikova M, Pinto C, Gomathinayagam S, Vujanovic M, et al. DNA2 drives processing and restart of reversed replication forks in human cells. J Cell Biol. 2015;208(5):545–62. doi: 10.1083/jcb.201406100. PubMed PMID: 25733713; PubMed Central PMCID: PMCPMC4347643.

26. Tye S, Ronson GE, Morris JR. A fork in the road: Where homologous recombination and stalled replication fork protection part ways. Semin Cell Dev Biol. 2021;113:14–26. Epub 20200709. doi: 10.1016/j.semcdb.2020.07.004. PubMed PMID: 32653304; PubMed Central PMCID: PMCPMC8082280.

27. Guh CL, Lei KH, Chen YA, Jiang YZ, Chang HY, Liaw H, et al. RAD51 paralogs synergize with RAD51 to protect reversed forks from cellular nucleases. Nucleic Acids Res. 2023;51(21):11717–31. doi: 10.1093/nar/gkad856. PubMed PMID: 37843130; PubMed Central PMCID: PMCPMC10681713.

28. Lemacon D, Jackson J, Quinet A, Brickner JR, Li S, Yazinski S, et al. MRE11 and EXO1 nucleases degrade reversed forks and elicit MUS81-dependent fork rescue in BRCA2-deficient cells. Nat Commun. 2017;8(1):860. Epub 20171016. doi: 10.1038/s41467-017-01180-5. PubMed PMID: 29038425; PubMed Central PMCID: PMCPMC5643552.

29. Sale JE. Translesion DNA synthesis and mutagenesis in eukaryotes. Cold Spring Harb Perspect Biol. 2013;5(3):a012708. Epub 20130301. doi: 10.1101/cshperspect.a012708. PubMed PMID: 23457261; PubMed Central PMCID: PMCPMC3578355.

30. Ghosal G, Chen J. DNA damage tolerance: a double-edged sword guarding the genome. Transl Cancer Res. 2013;2(3):107–29. doi: 10.3978/j.issn.2218-676X.2013.04.01. PubMed PMID: 24058901; PubMed Central PMCID: PMCPMC3779140.

31. Vaisman A, Woodgate R. Translesion DNA polymerases in eukaryotes: what makes them tick? Crit Rev Biochem Mol Biol. 2017;52(3):274–303. Epub 20170309. doi: 10.1080/10409238.2017.1291576. PubMed PMID: 28279077; PubMed Central PMCID: PMCPMC5573590.

32. Malik R, Kopylov M, Gomez-Llorente Y, Jain R, Johnson RE, Prakash L, et al. Structure and mechanism of B-family DNA polymerase zeta specialized for translesion DNA synthesis. Nat Struct Mol Biol. 2020;27(10):913–24. Epub 20200817. doi: 10.1038/s41594-020-0476-7. PubMed PMID: 32807989; PubMed Central PMCID: PMCPMC7554088.

33. Gomez-Llorente Y, Malik R, Jain R, Choudhury JR, Johnson RE, Prakash L, et al. The architecture of yeast DNA polymerase zeta. Cell Rep. 2013;5(1):79–86. Epub 20131010. doi: 10.1016/j.celrep.2013.08.046. PubMed PMID: 24120860; PubMed Central PMCID: PMCPMC3883112.

34. Makarova AV, Stodola JL, Burgers PM. A four-subunit DNA polymerase zeta complex containing Pol delta accessory subunits is essential for PCNA-mediated mutagenesis. Nucleic Acids Res. 2012;40(22):11618–26. Epub 20121012. doi: 10.1093/nar/gks948. PubMed PMID: 23066099; PubMed Central PMCID: PMCPMC3526297.

35. Netz DJ, Stith CM, Stumpfig M, Kopf G, Vogel D, Genau HM, et al. Eukaryotic DNA polymerases require an iron-sulfur cluster for the formation of active complexes. Nat Chem Biol. 2011;8(1):125–32. Epub 20111127. doi: 10.1038/nchembio.721. PubMed PMID: 22119860; PubMed Central PMCID: PMCPMC3241888.

36. Ma X, Tang TS, Guo C. Regulation of translesion DNA synthesis in mammalian cells. Environ Mol Mutagen. 2020;61(7):680–92. Epub 20200206. doi: 10.1002/em.22359. PubMed PMID: 31983077.

37. Davies AA, Huttner D, Daigaku Y, Chen S, Ulrich HD. Activation of ubiquitin-dependent DNA damage bypass is mediated by replication protein a. Mol Cell. 2008;29(5):625–36. doi: 10.1016/j.molcel.2007.12.016. PubMed PMID: 18342608; PubMed Central PMCID: PMCPMC2507760.

38. Hedglin M, Benkovic SJ. Regulation of Rad6/Rad18 Activity During DNA Damage Tolerance. Annu Rev Biophys. 2015;44:207–28. doi: 10.1146/annurev-biophys-060414-033841. PubMed PMID: 26098514; PubMed Central PMCID: PMCPMC5592839.

39. Andersen PL, Xu F, Xiao W. Eukaryotic DNA damage tolerance and translesion synthesis through covalent modifications of PCNA. Cell Res. 2008;18(1):162–73. doi: 10.1038/cr.2007.114. PubMed PMID: 18157158.

40. Boehm EM, Spies M, Washington MT. PCNA tool belts and polymerase bridges form during translesion synthesis. Nucleic Acids Res. 2016;44(17):8250–60. Epub 20160620. doi: 10.1093/nar/gkw563. PubMed PMID: 27325737; PubMed Central PMCID: PMCPMC5041468.

41. Edmunds CE, Simpson LJ, Sale JE. PCNA ubiquitination and REV1 define temporally distinct mechanisms for controlling translesion synthesis in the avian cell line DT40. Mol Cell. 2008;30(4):519–29. doi: 10.1016/j.molcel.2008.03.024. PubMed PMID: 18498753.

42. Livneh Z, Ziv O, Shachar S. Multiple two-polymerase mechanisms in mammalian translesion DNA synthesis. Cell Cycle. 2010;9(4):729–35. Epub 20100223. doi: 10.4161/cc.9.4.10727. PubMed PMID: 20139724.

43. Shachar S, Ziv O, Avkin S, Adar S, Wittschieben J, Reissner T, et al. Two-polymerase mechanisms dictate error-free and error-prone translesion DNA synthesis in mammals. EMBO J. 2009;28(4):383-93. Epub 20090115. doi: 10.1038/emboj.2008.281. PubMed PMID: 19153606; PubMed Central PMCID: PMCPMC2646147.

44. Sale JE, Lehmann AR, Woodgate R. Y-family DNA polymerases and their role in tolerance of cellular DNA damage. Nat Rev Mol Cell Biol. 2012;13(3):141–52. Epub 20120223. doi: 10.1038/nrm3289. PubMed PMID: 22358330; PubMed Central PMCID: PMCPMC3630503.

45. Wiltrout ME, Walker GC. The DNA polymerase activity of Saccharomyces cerevisiae Rev1 is biologically significant. Genetics. 2011;187(1):21–35. Epub 20101026. doi: 10.1534/genetics.110.124172. PubMed PMID: 20980236; PubMed Central PMCID: PMCPMC3018306.

46. Ross AL, Sale JE. The catalytic activity of REV1 is employed during immunoglobulin gene diversification in DT40. Mol Immunol. 2006;43(10):1587–94. Epub 20051102. doi: 10.1016/j.molimm.2005.09.017. PubMed PMID: 16263170.

47. Otsuka C, Loakes D, Negishi K. The role of deoxycytidyl transferase activity of yeast Rev1 protein in the bypass of abasic sites. Nucleic Acids Res Suppl. 2002;(2):87–8. doi: 10.1093/nass/2.1.87. PubMed PMID: 12903118.

48. Northam MR, Moore EA, Mertz TM, Binz SK, Stith CM, Stepchenkova EI, et al. DNA polymerases zeta and Rev1 mediate error-prone bypass of non-B DNA structures. Nucleic Acids Res. 2014;42(1):290–306. Epub 20130918. doi: 10.1093/nar/gkt830. PubMed PMID: 24049079; PubMed Central PMCID: PMCPMC3874155.

49. Sarkies P, Reams C, Simpson LJ, Sale JE. Epigenetic instability due to defective replication of structured DNA. Mol Cell. 2010;40(5):703–13. doi: 10.1016/j.molcel.2010.11.009. PubMed PMID: 21145480; PubMed Central PMCID: PMCPMC3145961.

50. Yang Y, Liu Z, Wang F, Temviriyanukul P, Ma X, Tu Y, et al. FANCD2 and REV1 cooperate in the protection of nascent DNA strands in response to replication stress. Nucleic Acids Res. 2015;43(17):8325–39. Epub 20150717. doi: 10.1093/nar/gkv737. PubMed PMID: 26187992; PubMed Central PMCID: PMCPMC4787816.

51. Kuang L, Kou H, Xie Z, Zhou Y, Feng X, Wang L, Wang Z. A non-catalytic function of Rev1 in translesion DNA synthesis and mutagenesis is mediated by its stable interaction with Rad5. DNA Repair (Amst). 2013;12(1):27–37. Epub 20121109. doi: 10.1016/j.dnarep.2012.10.003. PubMed PMID: 23142547.

52. Pages V, Bresson A, Acharya N, Prakash S, Fuchs RP, Prakash L. Requirement of Rad5 for DNA polymerase zeta-dependent translesion synthesis in Saccharomyces cerevisiae. Genetics. 2008;180(1):73–82. Epub 20080830. doi: 10.1534/genetics.108.091066. PubMed PMID: 18757916; PubMed Central PMCID: PMCPMC2535723.

53. Kane DP, Shusterman M, Rong Y, McVey M. Competition between replicative and translesion polymerases during homologous recombination repair in Drosophila. PLoS Genet. 2012;8(4):e1002659. doi: 10.1371/journal.pgen.1002659. PubMed PMID: 22532806; PubMed Central PMCID: PMCPMC3330096.

54. Zamborszky J, Szikriszt B, Gervai JZ, Pipek O, Poti A, Krzystanek M, et al. Loss of BRCA1 or BRCA2 markedly increases the rate of base substitution mutagenesis and has distinct effects on genomic deletions. Oncogene. 2017;36(35):5085–6. Epub 20170626. doi: 10.1038/onc.2017.213. PubMed PMID: 28650471; PubMed Central PMCID: PMCPMC5582208.

55. Rosenbaum JC, Bonilla B, Hengel SR, Mertz TM, Herken BW, Kazemier HG, et al. The Rad51 paralogs facilitate a novel DNA strand specific damage tolerance pathway. Nat Commun. 2019;10(1):3515. Epub 20190805. doi: 10.1038/s41467-019-11374-8. PubMed PMID: 31383866; PubMed Central PMCID: PMCPMC6683157.

56. Conde F, San-Segundo PA. Role of Dot1 in the response to alkylating DNA damage in Saccharomyces cerevisiae: regulation of DNA damage tolerance by the error-prone polymerases Polzeta/Rev1. Genetics. 2008;179(3):1197–210. doi: 10.1534/genetics.108.089003. PubMed PMID: 18562671; PubMed Central PMCID: PMCPMC2475726.

57. McVey M, Adams M, Staeva-Vieira E, Sekelsky JJ. Evidence for multiple cycles of strand invasion during repair of double-strand gaps in Drosophila. Genetics. 2004;167(2):699–705. doi: 10.1534/genetics.103.025411. PubMed PMID: 15238522; PubMed Central PMCID: PMCPMC1470890.

58. Thomas AM, Hui C, South A, McVey M. Common variants of Drosophila melanogaster Cyp6d2 cause camptothecin sensitivity and synergize with loss of Brca2. G3 (Bethesda). 2013;3(1):91-9. Epub 20130101. doi: 10.1534/g3.112.003996. PubMed PMID: 23316441; PubMed Central PMCID: PMCPMC3538347.

59. Khodaverdian VY, McVey M. Rapid Detection of gamma-H2Av Foci in Ex Vivo MMS-Treated Drosophila Imaginal Discs. Methods Mol Biol. 2017;1644:203–11. doi: 10.1007/978-1-4939-7187-9_19. PubMed PMID: 28710767.

60. Guo C, Fischhaber PL, Luk-Paszyc MJ, Masuda Y, Zhou J, Kamiya K, et al. Mouse Rev1 protein interacts with multiple DNA polymerases involved in translesion DNA synthesis. EMBO J. 2003;22(24):6621–30. doi: 10.1093/emboj/cdg626. PubMed PMID: 14657033; PubMed Central PMCID: PMCPMC291821.

61. Tissier A, Kannouche P, Reck MP, Lehmann AR, Fuchs RP, Cordonnier A. Co-localization in replication foci and interaction of human Y-family members, DNA polymerase pol eta and REVl protein. DNA Repair (Amst). 2004;3(11):1503–14. doi: 10.1016/j.dnarep.2004.06.015. PubMed PMID: 15380106.

62. Ohashi E, Murakumo Y, Kanjo N, Akagi J, Masutani C, Hanaoka F, Ohmori H. Interaction of hREV1 with three human Y-family DNA polymerases. Genes Cells. 2004;9(6):523–31. doi: 10.1111/j.1356-9597.2004.00747.x. PubMed PMID: 15189446.

63. Acharya N, Haracska L, Johnson RE, Unk I, Prakash S, Prakash L. Complex formation of yeast Rev1 and Rev7 proteins: a novel role for the polymerase-associated domain. Mol Cell Biol. 2005;25(21):9734–40. doi: 10.1128/MCB.25.21.9734-9740.2005. PubMed PMID: 16227619; PubMed Central PMCID: PMCPMC1265840.

64. Haracska L, Acharya N, Unk I, Johnson RE, Hurwitz J, Prakash L, Prakash S. A single domain in human DNA polymerase iota mediates interaction with PCNA: implications for translesion DNA synthesis. Mol Cell Biol. 2005;25(3):1183–90. doi: 10.1128/MCB.25.3.1183-1190.2005. PubMed PMID: 15657443; PubMed Central PMCID: PMCPMC544020.

65. Kosarek JN, Woodruff RV, Rivera-Begeman A, Guo C, D’Souza S, Koonin EV, et al. Comparative analysis of in vivo interactions between Rev1 protein and other Y-family DNA polymerases in animals and yeasts. DNA Repair (Amst). 2008;7(3):439–51. doi: 10.1016/j.dnarep.2007.11.016. PubMed PMID: 18242152; PubMed Central PMCID: PMCPMC2363158.

66. Gao G, McMahon C, Chen J, Rong YS. A powerful method combining homologous recombination and site-specific recombination for targeted mutagenesis in Drosophila. Proc Natl Acad Sci U S A. 2008;105(37):13999–4004. doi: 10.1073/pnas.0805843105. PubMed PMID: 18772376; PubMed Central PMCID: PMCPMC2529331.

67. Gao G, Wesolowska N, Rong YS. SIRT combines homologous recombination, site-specific integration, and bacterial recombineering for targeted mutagenesis in Drosophila. Cold Spring Harb Protoc. 2009;2009(6):pdb prot5236. doi: 10.1101/pdb.prot5236. PubMed PMID: 20147194.

68. Zhang Y, Schreiner W, Rong YS. Genome manipulations with bacterial recombineering and site-specific integration in Drosophila. Methods Mol Biol. 2014;1114:11–24. doi: 10.1007/978-1-62703-761-7_2. PubMed PMID: 24557894.

69. Wojtaszek J, Lee CJ, D’Souza S, Minesinger B, Kim H, D’Andrea AD, et al. Structural basis of Rev1-mediated assembly of a quaternary vertebrate translesion polymerase complex consisting of Rev1, heterodimeric polymerase (Pol) zeta, and Pol kappa. J Biol Chem. 2012;287(40):33836-46. Epub 20120802. doi: 10.1074/jbc.M112.394841. PubMed PMID: 22859295; PubMed Central PMCID: PMCPMC3460478.

70. Bienko M, Green CM, Crosetto N, Rudolf F, Zapart G, Coull B, et al. Ubiquitin-binding domains in Y-family polymerases regulate translesion synthesis. Science. 2005;310(5755):1821-4. doi: 10.1126/science.1120615. PubMed PMID: 16357261.

71. Kikuchi S, Hara K, Shimizu T, Sato M, Hashimoto H. Crystallization and X-ray diffraction analysis of the ternary complex of the C-terminal domain of human REV1 in complex with REV7 bound to a REV3 fragment involved in translesion DNA synthesis. Acta Crystallogr Sect F Struct Biol Cryst Commun. 2012;68(Pt 8):962–4. Epub 20120727. doi: 10.1107/S1744309112032435. PubMed PMID: 22869133; PubMed Central PMCID: PMCPMC3412784.

72. McPherson KS, Rizzo AA, Erlandsen H, Chatterjee N, Walker GC, Korzhnev DM. Evolution of Rev7 interactions in eukaryotic TLS DNA polymerase Polzeta. J Biol Chem. 2023;299(2):102859. Epub 20221231. doi: 10.1016/j.jbc.2022.102859. PubMed PMID: 36592930; PubMed Central PMCID: PMCPMC9926120.

73. Guo C, Sonoda E, Tang TS, Parker JL, Bielen AB, Takeda S, et al. REV1 protein interacts with PCNA: significance of the REV1 BRCT domain in vitro and in vivo. Mol Cell. 2006;23(2):265–71. doi: 10.1016/j.molcel.2006.05.038. PubMed PMID: 16857592.

74. Pustovalova Y, Maciejewski MW, Korzhnev DM. NMR mapping of PCNA interaction with translesion synthesis DNA polymerase Rev1 mediated by Rev1-BRCT domain. J Mol Biol. 2013;425(17):3091–105. Epub 20130607. doi: 10.1016/j.jmb.2013.05.029. PubMed PMID: 23747975.

75. de Groote FH, Jansen JG, Masuda Y, Shah DM, Kamiya K, de Wind N, Siegal G. The Rev1 translesion synthesis polymerase has multiple distinct DNA binding modes. DNA Repair (Amst). 2011;10(9):915–25. Epub 20110712. doi: 10.1016/j.dnarep.2011.04.033. PubMed PMID: 21752727.

76. Bomar MG, D’Souza S, Bienko M, Dikic I, Walker GC, Zhou P. Unconventional ubiquitin recognition by the ubiquitin-binding motif within the Y family DNA polymerases iota and Rev1. Mol Cell. 2010;37(3):408–17. doi: 10.1016/j.molcel.2009.12.038. PubMed PMID: 20159559; PubMed Central PMCID: PMCPMC2841503.

77. Niu X, Chen W, Bi T, Lu M, Qin Z, Xiao W. Rev1 plays central roles in mammalian DNA-damage tolerance in response to UV irradiation. FEBS J. 2019;286(14):2711–25. Epub 20190411. doi: 10.1111/febs.14840. PubMed PMID: 30963698.

78. Jansen JG, Tsaalbi-Shtylik A, Langerak P, Calleja F, Meijers CM, Jacobs H, de Wind N. The BRCT domain of mammalian Rev1 is involved in regulating DNA translesion synthesis. Nucleic Acids Res. 2005;33(1):356–65. Epub 20050113. doi: 10.1093/nar/gki189. PubMed PMID: 15653636; PubMed Central PMCID: PMCPMC546167.

79. Sasatani M, Zaharieva EK, Kamiya K. The in vivo role of Rev1 in mutagenesis and carcinogenesis. Genes Environ. 2020;42:9. Epub 20200228. doi: 10.1186/s41021-020-0148-1. PubMed PMID: 32161626; PubMed Central PMCID: PMCPMC7048032.

80. Zhou Y, Wang J, Zhang Y, Wang Z. The catalytic function of the Rev1 dCMP transferase is required in a lesion-specific manner for translesion synthesis and base damage-induced mutagenesis. Nucleic Acids Res. 2010;38(15):5036–46. Epub 20100412. doi: 10.1093/nar/gkq225. PubMed PMID: 20388628; PubMed Central PMCID: PMCPMC2926598.

81. Guo C, Tang TS, Bienko M, Parker JL, Bielen AB, Sonoda E, et al. Ubiquitin-binding motifs in REV1 protein are required for its role in the tolerance of DNA damage. Mol Cell Biol. 2006;26(23):8892–900. Epub 20060918. doi: 10.1128/MCB.01118-06. PubMed PMID: 16982685; PubMed Central PMCID: PMCPMC1636806.

82. Masuda Y, Kamiya K. Role of single-stranded DNA in targeting REV1 to primer termini. J Biol Chem. 2006;281(34):24314–21. Epub 20060627. doi: 10.1074/jbc.M602967200. PubMed PMID: 16803901.

83. Zhang Y, Wu X, Rechkoblit O, Geacintov NE, Taylor JS, Wang Z. Response of human REV1 to different DNA damage: preferential dCMP insertion opposite the lesion. Nucleic Acids Res. 2002;30(7):1630–8. doi: 10.1093/nar/30.7.1630. PubMed PMID: 11917024; PubMed Central PMCID: PMCPMC101843.

84. Stone JE, Kumar D, Binz SK, Inase A, Iwai S, Chabes A, et al. Lesion bypass by S. cerevisiae Pol zeta alone. DNA Repair (Amst). 2011;10(8):826–34. Epub 20110531. doi: 10.1016/j.dnarep.2011.04.032. PubMed PMID: 21622032; PubMed Central PMCID: PMCPMC3146559.

85. Korzhnev DM, Hadden MK. Targeting the Translesion Synthesis Pathway for the Development of Anti-Cancer Chemotherapeutics. J Med Chem. 2016;59(20):9321–36. Epub 20160719. doi: 10.1021/acs.jmedchem.6b00596. PubMed PMID: 27362876.

86. Russo M, Crisafulli G, Sogari A, Reilly NM, Arena S, Lamba S, et al. Adaptive mutability of colorectal cancers in response to targeted therapies. Science. 2019;366(6472):1473-80. Epub 20191107. doi: 10.1126/science.aav4474. PubMed PMID: 31699882.

87. Temprine K, Campbell NR, Huang R, Langdon EM, Simon-Vermot T, Mehta K, et al. Regulation of the error-prone DNA polymerase Polkappa by oncogenic signaling and its contribution to drug resistance. Sci Signal. 2020;13(629). Epub 20200428. doi: 10.1126/scisignal.aau1453. PubMed PMID: 32345725; PubMed Central PMCID: PMCPMC7428051.

88. Xie K, Doles J, Hemann MT, Walker GC. Error-prone translesion synthesis mediates acquired chemoresistance. Proc Natl Acad Sci U S A. 2010;107(48):20792–7. Epub 20101110. doi: 10.1073/pnas.1011412107. PubMed PMID: 21068378; PubMed Central PMCID: PMCPMC2996453.

89. Wojtaszek JL, Chatterjee N, Najeeb J, Ramos A, Lee M, Bian K, et al. A Small Molecule Targeting Mutagenic Translesion Synthesis Improves Chemotherapy. Cell. 2019;178(1):152–9 e11. Epub 20190606. doi: 10.1016/j.cell.2019.05.028. PubMed PMID: 31178121; PubMed Central PMCID: PMCPMC6644000.

90. Chatterjee N, Whitman MA, Harris CA, Min SM, Jonas O, Lien EC, et al. REV1 inhibitor JH-RE-06 enhances tumor cell response to chemotherapy by triggering senescence hallmarks. Proc Natl Acad Sci U S A. 2020;117(46):28918–21. Epub 20201109. doi: 10.1073/pnas.2016064117. PubMed PMID: 33168727; PubMed Central PMCID: PMCPMC7682577.

91. Staeva-Vieira E, Yoo S, Lehmann R. An essential role of DmRad51/SpnA in DNA repair and meiotic checkpoint control. EMBO J. 2003;22(21):5863–74. doi: 10.1093/emboj/cdg564. PubMed PMID: 14592983; PubMed Central PMCID: PMCPMC275421.

92. Klovstad M, Abdu U, Schupbach T. Drosophila brca2 is required for mitotic and meiotic DNA repair and efficient activation of the meiotic recombination checkpoint. PLoS Genet. 2008;4(2):e31. doi: 10.1371/journal.pgen.0040031. PubMed PMID: 18266476; PubMed Central PMCID: PMCPMC2233675.

93. Adler PN, MacQueen M. Cell proliferation and DNA replication in the imaginal wing disc of Drosophila melanogaster. Dev Biol. 1984;103(1):28–37. doi: 10.1016/0012-1606(84)90004-6. PubMed PMID: 6425098.

94. Gatti M, Santini G, Pimpinelli S, Olivieri G. Lack of spontaneous sister chromatid exchanges in somatic cells of Drosophila melanogaster. Genetics. 1979;91(2):255–74. doi: 10.1093/genetics/91.2.255. PubMed PMID: 109350; PubMed Central PMCID: PMCPMC1216365.

